# Tinamou egg color displacement at ecoregion co-partitioning

**DOI:** 10.1101/2020.04.26.062927

**Authors:** Qin Li, Silu Wang

## Abstract

The divergence of reproductive traits frequently underpins the evolution of reproductive isolation. One of the most enduring puzzles on this subject concerns the variability in egg coloration among species of tinamou (Tinamidae), endemic to neotropics. Here we investigated the hypothesis that tinamou egg coloration is a mating signal and its diversification was driven by reinforcement. For most tinamou species, the male guards the nest that is sequentially visited and laid eggs in by multiple females. The colorations of the existing eggs in the nest could signal mate quality and species identities to the upcoming females, preventing costly hybridization, thus were selected to diverge among species (Mating Signal Character Displacement Hypothesis). If so, two predictions should follow: (1) egg colors should coevolve with known mating signals as the tinamou lineages diverged; (2) species that partition similar ecoregions should display different egg colors. The tinamou songs are important mating signals and are highly divergent among species. We found that the egg luminance was significantly associated with the first principal component of the song variables, which supports prediction (1). In addition, we found support for (2): tinamou species that co-partition ecoregions tend to display different egg colors, controlling for song variation. Egg color and songs could be multimodal mating signals that are divergently selected as different tinamou species diverged. Mating signal evolution could be opportunistic and even exploit post-mating trait as premating signals that undergo character displacement at sympatry.

## Introduction

Reproductive trait divergence is crucial for speciation because these traits frequently underpin barriers to gene flow in the onset of speciation (Koski and Ashman 2016; Pfennig 2016). A very puzzling reproductive trait divergence exists in tinamou egg coloration. Tinamiformes (common name: tinamou) is the most species-rich order of Palaeognathae, containing 48 extant species (Figure 1) (Cabot 1992; Davies 2002). Tinamous are mostly dull in plumage, but various species of tinamous lay brightly colored eggs ranging from brilliant magenta, to pink, purple, turquoise, and olive green (Figure 1) (Cabot 1992). Despite its evolutionary uniqueness, tinamou egg color divergence remains a mystery due to the cryptic and elusive plumage and behavior of the birds (Cabot 1992; Davies 2002).

**Figure 1.**
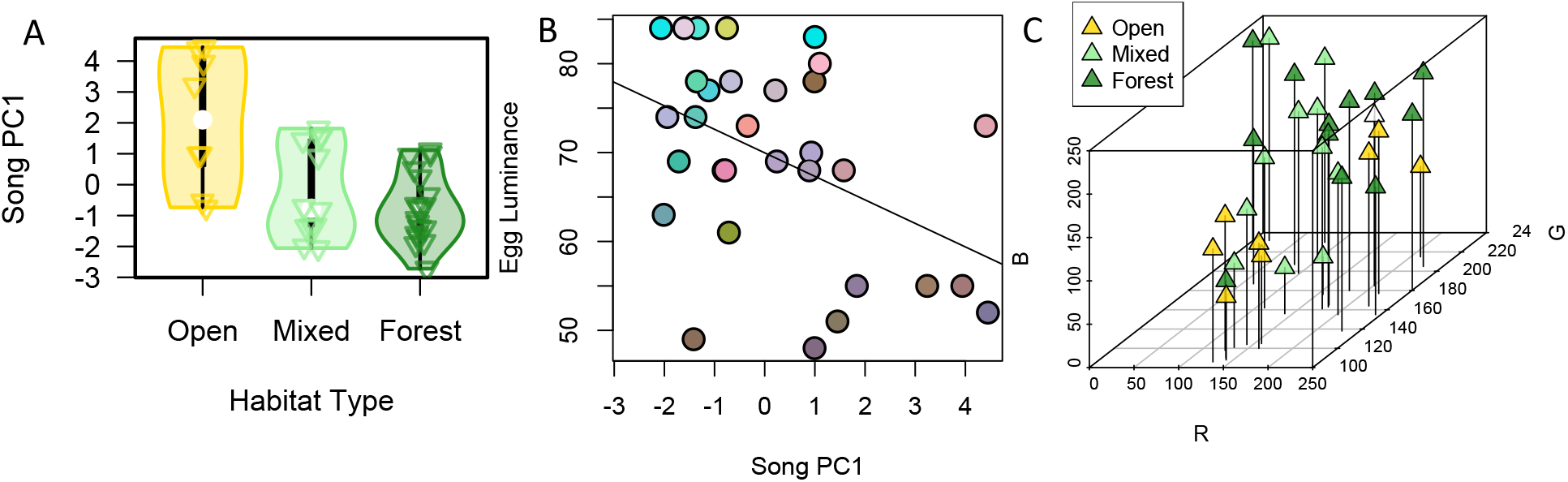
Association of habitat type, song variation, and egg colour among tinamou species. (A) There is an association between song variation and habitat type among tinamou species (Bertelli and Tubaro 2002). (B) There is significant association between egg luminance and song PC1 (permutation test of independence, Z = -2.42, R^2^ = 0.20, *p* = 0.015, *p*_*corrected*_ = 0.045). The color of dots represent the RGB color of the eggs of each species. (C) Egg color PC1 in the RGB space is not significantly associated with habitat types among tinamou species (*p* > 0.05).

Two main hypotheses have been made about evolutionary mechanism of egg color divergence: (1) ‘aposematism hypothesis’ (Swynnerton 1916) states that eggs are brightly colored to warn predators of their distastefulness; (2) ‘mating signaling character displacement hypothesis’, which predicts that egg color are employed for mating recognition and attraction (Weeks 1973; Brennan 2010), and could be divergently selected among closely related species at sympatry. The aposematic function of egg coloration was thought to be unlikely, because egg predators are nocturnal mammals or reptiles that prioritize chemical cues over visual cues (Skutch 1966; Cabot 1992; Brennan 2010). The ‘mating signal character displacement hypothesis’ is plausible considering the ecology and mating systems of tinamou species. In most tinamou species, males collect and incubate eggs laid by multiple females (Cabot 1992; Davies 2002). An initial female is attracted by distinctive songs to the nest guarded a male. If mating occurs, the female lays eggs in the nest and leaves before other females come and lay more eggs that are all incubated by the same male (Cabot 1992; Davies 2002). The colors of the existing eggs in the nest could signal intra-specific mate identity to prevent costly hybridization. Such alternative “mating channel” might be especially favored when the plumage is adapted for camouflage (Cabot 1992; Davies 2002). If egg colors of different tinamou species is adapted for mate recognition, reinforcement (Dobzhansky 1937; Mayr 1942; Liou and Price 1994; Servedio 2000) could have driven egg color divergence among different tinamou species.

Here we investigated the ‘mating signal character displacement hypothesis’ of tinamou egg color divergence. If tinamou egg colors are employed as a species-specific mate recognition signal, it should coevolve with other mate-recognition signals as the tinamou lineages diverged. Songs of tinamous are known to be important and highly divergent mating signals among species (Cabot 1992; Bertelli and Tubaro 2002; Laverde-R. and Cadena 2014; Boesman et al. 2018). In addition to songs, multimodality of mating signals is selected for efficient mate-recognition in complex environment (Partan and Marler 1999; Rowe 1999; Secondi et al. 2015; Halfwerk et al. 2019). When glamorous plumage is costly due to increased predation risk, alternative signal modality such as egg coloration could be favored. Since the birds themselves (instead of the eggs) tend to attract nest predation (Brennan 2010), egg coloration could be employed to fulfill signal multimodality, bypassing the evolutionary constraints for plumage. To investigate this possibility, here we ask: (1) whether egg coloration coevolved with songs as tinamou species diverged; and (2) whether species that co-partition ecoregions tend to display divergent egg colors?

## Methods

To investigate the mating signal character displacement hypothesis in tinamou egg color evolution, we examined the association of song, egg colors, and ecoregion co-partitions among tinamou species. Tinamou egg coloration data was extracted from existing tinamou egg color analysis (Schläpfer 2017), which quantified coloration of the eggs from 32 tinamou species. Since tinamou egg colors are known to decay over the course museum storage, tinamou nest photos are used to best represent the functional egg color (Schläpfer 2017). Briefly, for each species, the egg coloration reflected in hue, chroma, and luminance in the CIELAB color space as well as RGB color space was quantified (Schläpfer 2017). To efficiently infer egg color variation among species, we conducted principal component analysis (PCA) of the RGB color space with RGB axes shifted to center around zero and scaled to unit variance.

Song data was acquired from a previous study (Bertelli and Tubaro 2002), in which four song variables from 40 tinamou species were quantified: maximum frequency (Hz), minimum frequency (Hz), emphasized frequency (frequency of the note with highest amplitude in the song, Hz), and bandwidth (the difference between maximum and minimum frequency, Hz). Because these variables are correlated (Figure S1), we used PCA to generate PC1 that captured 81.2% of the variation in song data. Four song variables were shifted to center around zero and scaled to unit variance preceding PCA.

We tested whether egg hue, chroma, and luminance is respectively associated with song PC1 among tinamous, accounting for the potential confounding phylogenetic signals as well as habitat type, which was shown to be correlated with song variation among tinamous (Figure 1 A)(Bertelli and Tubaro 2002). We used the Tinamidae phylogenetic tree inferred with both molecular (1143 bp mitochondrial and 1145 bp nuclear) and morphological characters (237 characters) (Bertelli and Porzecanski 2004), in which the polytomy was resolved with additional morphological and life history traits (Bertelli 2017). We first tested if there is phylogenetic signal (Pagel 1999) in each of the egg color variable with phytools (Revell 2012). If no significant phylogenetic signal was identified, we ran permutation test of independence between song PC1 and the egg color variable with *coin* package. If there is significant (*p* < 0.05) phylogenetic signal, we first calculated independent contrasts of egg colors and song PC1 with Ape (Paradis and Schliep 2019), then executed independence test. To correct for multiple hypotheses testing, we conducted False Discovery Rate correction (Benjamini and Hochberg 1995) to correct for three tests.

Species ranges were generated according to their occurrences. Geo-referenced occurrences were downloaded from GBIF (Global Biodiversity Information Facility, GBIF.org) for all the tinamou species. Occurrences with uncertainty in coordinates less than 10km were kept, and were further filtered by removing redundant localities within a 1-km^2^ grid cell via R package *ecospat* (Cola et al. 2017) after Lambert azimuthal equal-area projection (ranging 9 – 9413, median = 469). Ecoregion designation for each species was carried out by overlaying occurrences with the ecoregion map of America. Ecoregion maps were downloaded for North America (https://www.epa.gov) and South America (http://ecologicalregions.info) separately at level I and then were combined as one to include 11 ecoregions for all occurrences. For each species, an ecoregion with which the proportion of occurrences was greater than 10% (or > 50 if the total number > 1000) was included in the species’ ecoregion distribution.

With this data, we calculated pairwise probability of ecoregion co-partitioning among tinamou species. For each pair of species, the probability of ecoregion co-partitioning is the sum (across all ecoregions) of the multiplication of the probability of species-specific partitioning within each ecoregion. For each species, the probability of partitioning within an ecoregion is the number of points of occurrence within the ecoregion over the total points of occurrence for the species. If the probability of ecoregion co-partitioning is an effective indicator of sympatry or parapatry in the course of tinamou speciation, closely related species should be more likely to co-partition ecoregions. We employed Mantel Pearson’s correlation test with function *mantel*.*test* to test if the species distance matrix is correlated with the ecoregion co-partitioning probability matrix.

Further, we examined the evolutionary relationship between the PC1 of the egg RGB color space and ecoregion co-partitioning while controlling for song PC1 among species. We first computed the distance matrices of egg color PC1 and song PC1 respectively with the *dist* function in R. To examined matrix correlation between the distance matrix of egg color PC1 and the probability matrix of ecoregion co-partitioning while controlling for song PC1 distance matrix among species, we used one-tailed Partial Mantel Pearson’s correlation test with 10,000 iterations with the *mantel*.*partial* function.

## Results

The macroevolutionary relation among egg color, ecoregion partitioning, and song characters is consistent with the Character Displacement Hypothesis. We found a significant association between tinamou songs and egg luminance. Song PC1 represents 81.2% variation among the song variables in tinamous. Significant phylogenetic signal was observed in egg hue (K = 0.43, *p* = 0.049), chroma (K= 0.96, *p* = 0.001), and habitat type (K = 1.15, *p* = 0.001), but not in luminance (K = 0.37, *p* = 0.17). The tinamou song PC1 was significantly associated with egg luminance (Z = -2.42, *p* = 0.015, *p*_*corrected*_ = 0.045; Figure 1B), but not with egg hue (Z = 0.34, *p* = 0.72, *p*_*corrected*_ = 0.75), nor with egg chroma (Z = -0.32, *p* = 0.75, *p*_*corrected*_ = 0.75). The habitat type was although associated with tinamou song variation (Bertelli and Tubaro 2002), not significantly associated with egg color (Figure 1C): luminance (Phylogenetic ANOVA, F = 1.18, *p* = 0.82), chroma (F = 0.99, *p* = 0.84), or hue (F = 3.58, *p* = 0.53).

We further observed greater egg color divergence between species that co-partition ecoregions after controlling for song variation among tinamou species. The PC1 explains 54% variation of egg colors in the RGB color space. The distance of egg color PC1 distance is significantly positively associated with the ecoregion co-partitioning among species controlling for song distance among tinamou species (Partial Mantel r = 0.066, *p* = 0.048). Ecoregion co-occurrence is likely an effective indicator of parapatry or sympatry in history of tinamou speciation, as closely related species tends to co-occur at ecoregions (Mantel Z = 1530.38, *p* = 0.001).

## Discussion

The macroevolution relationship among egg colors, song variation, and ecoregion partitioning, is consistent with the ‘Mating Signal Character Displacement Hypothesis’ that tinamou egg coloration is an alternative mating signal that is divergently selected among species with similar ecogeographical partition (Weeks 1973; Brennan 2010). Egg colors and songs could be divergently selected as multimodal mate-recognition signals among tinamou species with similar appearance that partition similar ecogeographical range. When the plumage modality is constrained by anti-predation adaptation, egg coloration could be opportunistically adopted to fulfill mating signal multimodality for species recognition at sympatry. This study sheds light on the evolution of multimodal sexual signals that bypasses natural selection for plumage camouflage in the most species-rich order of Palaeognathae.

Egg coloration could be both a pre- and post-mating signals in tinamous. In many other bird species, egg color is known to be post-mating signals indicating female quality and to influence male incubation and promiscuity (Soler et al. 2005; English 2009). In tinamous, egg coloration can also function as premating signal as well, received by females. For most of the tinamou species, males guard and incubate the eggs laid by multiple females (Cabot 1992). The coloration of the existing eggs in the nest could be a mating signal received by the upcoming females to the nest. As premating signals, egg colors could reflect mate species identity (Sætre et al. 1997; Servedio and Noor 2003; Secondi et al. 2015) and mate quality, and stimulate female mate choice copying (Dugatkin 1992; Gibson and Höglund 1992).

Then why do tinamous adopt egg colors as mating signals in addition to songs? Songs that are important mating signals in tinamous (Bertelli and Tubaro 2002) and are involved in duetting between mating partners (Boesman et al. 2018). However, such acoustic modality is usually insufficient for mate communication in complex environment, thus signal redundancy or multimodality is further selected for mate communication to alleviate mate-searching effort and/or hybridization (Partan and Marler 1999; Rowe 1999; Uy et al. 2008; Secondi et al. 2015). Plumage patterning and coloration are frequently adopted as inter-specific and intra-specific mating signals at a finer spatial scale in birds (Uy et al. 2008, 2009; Seddon et al. 2013). However, most tinamou species exhibited cryptic plumage as an adaption for anti-predatory camouflage (Cabot 1992; Davies 2002). Sexual selection for plumage elaboration would compromise the adaptation for camouflage favored by natural selection. Egg coloration can be an alternative modality of mate signaling enrichment towards mate-searching refinement, bypassing the conflict between natural and sexual selection.

Reinforcement may have driven egg color and song divergence among sympatric or parapatric tinamou species. Many closely-related tinamou species demonstrate historical and/or contemporary ecoregion overlap (Cabot 1992)(Fig. 2), suggesting sympatry in the course of their speciation history. Ecological and/or intrinsic incompatibility among diverged lineages leads to reduced hybrid fitness, which in turn select for premating signal divergence to avoid costly hybridization (Dobzhansky 1940; Mayr 1942; Liou and Price 1994; Servedio 2000). Mating signal multimodality is needed to ensure mate recognition in complex heterospecific environment (Uy et al. 2008; Secondi et al. 2015). The divergence of tinamou song and egg coloration could jointly reduce heterospecific reproductive efforts among sympatric tinamou species. Closely related tinamou species that are mostly allopatric at present, but could still harbor footprints of historical character displacement of egg colors formed at historical sympatry either over secondary contact or sympatry speciation. Notably, the species with greater likelihood of ecoregion co-occurrence tend to display divergent egg colors (Fig. 2). For example, the closely related species *Crypturellus erythropus, C. notivagus, C. variegatus, C. brevirostris* still partition similar ecoregions and demonstrate distinct egg coloration, which may have been driven by reinforcement at historical sympatry (Fig. 3). Although the speciation event among tinamou species have long passed (∼34% sequence divergence), the likelihood of ecoregion co-occurrence is potentially an effective indicator of historical sympatry/parapatry in the history of tinamou speciation because the closely related species tend to share ecoregions.

**Figure 2.**
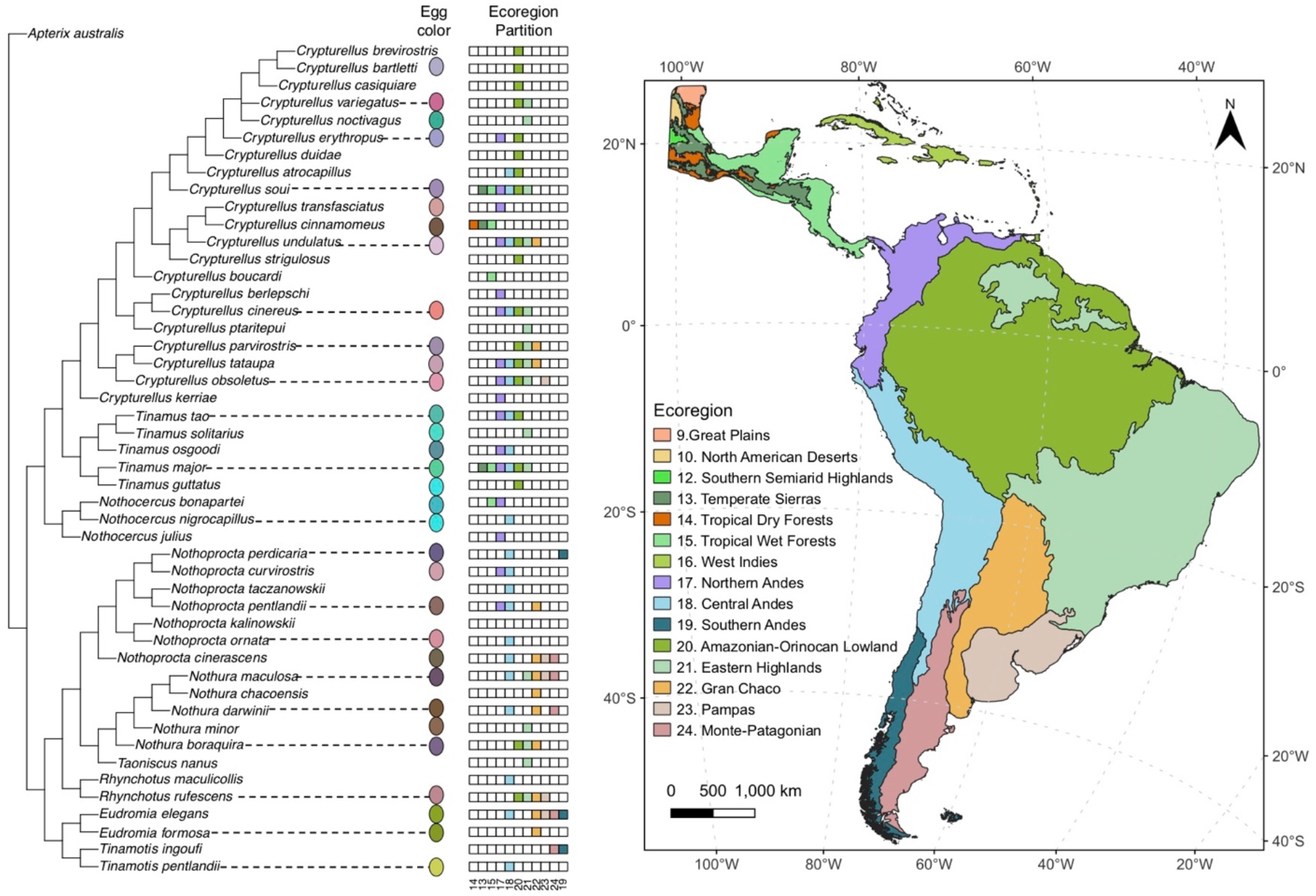
Tinamous ecoregion co-partitioning is associated with egg colour difference among species. There is significantly greater distance in egg colour among species that partition similar ecoregions, after controlling for song variation (partial mantel test, *p* < 0.05). The Tinamidae phylogeny was inferred with molecular and morphological data (Bertelli and Porzecanski 2004).

**Figure 3.**
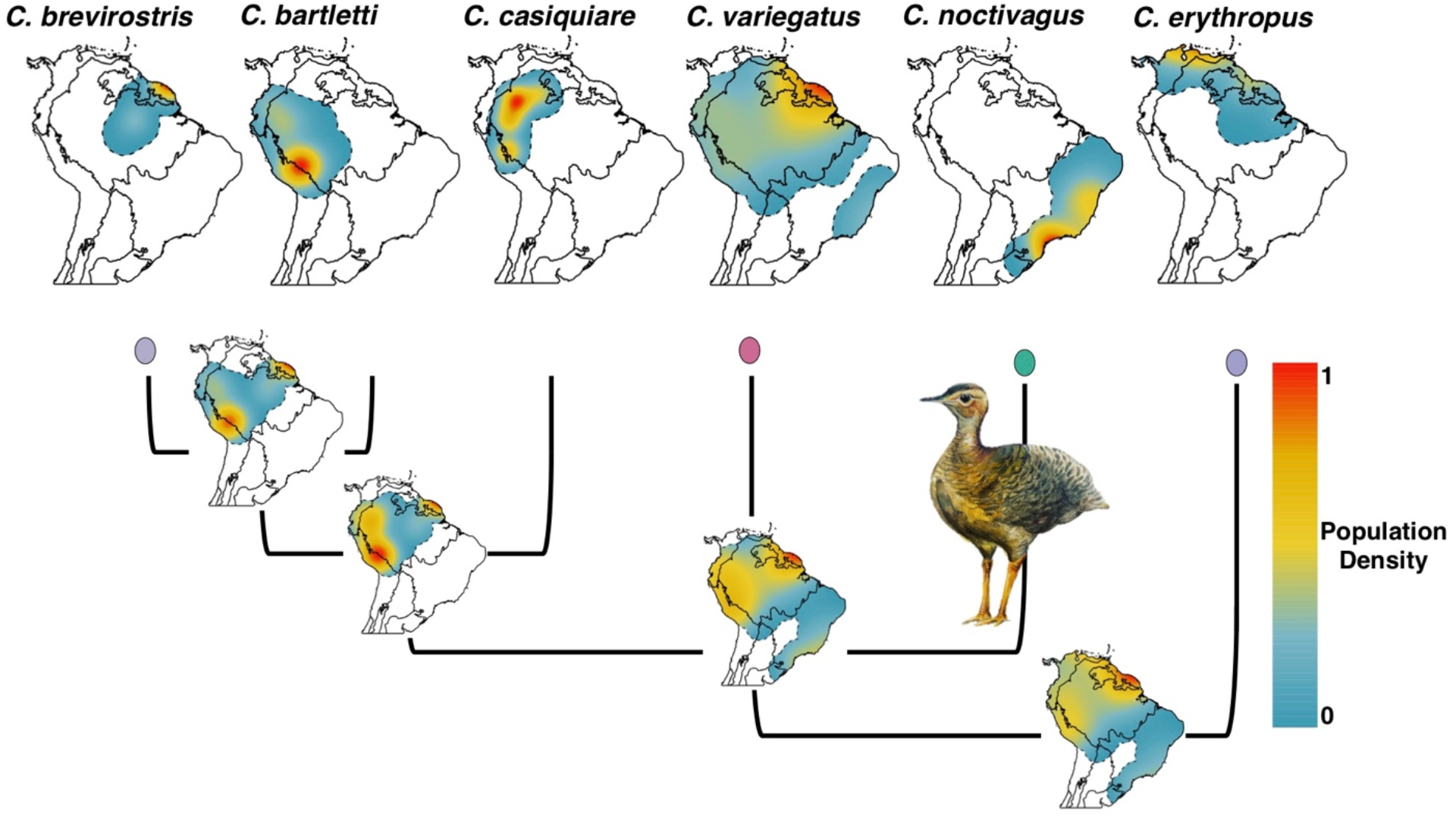
Reconstructed ancestral distribution of a *Crypturellus* clade. Black lines depict the boundaries of ecoregions corresponding to Figure 2, and dashed lines depicted boundaries of species’ distribution ranges. Ranges of tips were concave hull polygons generated via R package *rangemap* (Cobos et al. 2021) with a hierarchical clustering method, and kernel densities were estimated via R package *spatstat* (Adrian Baddeley, Ege Rubak 2015), to capture the spatial configuration of the occurrences. Distributions of internal nodes were generated by the same method with an intermediate step of rescaling densities (at a 100-km-resolution) of all descendants to offset the effect of variation in abundance. Blue to red colors depict the relative scale of kernel densities from low to high (ranging 0 to 1).

Although we found significant supports for both of the predictions of the Mating Signal Character Displacement Hypothesis, the significance levels were marginal. This indicates that there are other potential evolutionary forces shaping tinamou egg color divergence. Besides the local tinamou species assembly, other complex aspects of biotic interactions such as sexual conflicts, predation, and parasitism, might contribute to the egg color signal diversity.

The genetic underpinning of such song and egg color association is unclear. The simplest genetic mechanism underlying such association is pleiotropy (one gene affecting multiple traits) (Fisher 1930; Williams 1957). For example, the pleiotropic *foraging* gene underpins the association among social behavior and life history traits in natural populations (de Belle et al. 1989; Mery et al. 2007; Wang and Sokolowski 2017). There might be an omnipotent pleiotropic gene linking egg coloration, songs, among many other traits. If so, a wide array of traits underpinned by such pleiotropy are expected to coevolve. However, the association between song and egg color of tinamous is fairly specific: only luminance (but not hue or chroma) of the egg color is associated with tinamou songs. Such specificity in song-egg-color association is consistent with mating signal multimodality in which specifically signals were coupled among modalities (Gilliard 1956; Partan and Marler 1999; Hebets and Papaj 2005). Various signal modality can be underpinned by different genes and selected to coevolve as multimodal signals. Future investigation of genetic underpinnings of tinamou song and egg colors would further shed light on this interesting co-divergence. In sum, the results presented herein are concordant with the hypothesis that egg coloration in tinamous serves as mating signals either for mate attraction or mate species-recognition and could be divergently selected upon historical sympatry/parapatry.

## Acknowledgement

Thank Dahong Chen, Patricia Brennan, Daniel Hanley, Julia Clarke, and Jonathan Rolland for helpful discussions. Thank Daniel Hanley, Ken Thompson, Dahong Chen, and Jonathan Rolland for comments on the manuscript.

## Supplementary information

**Figure S1.**
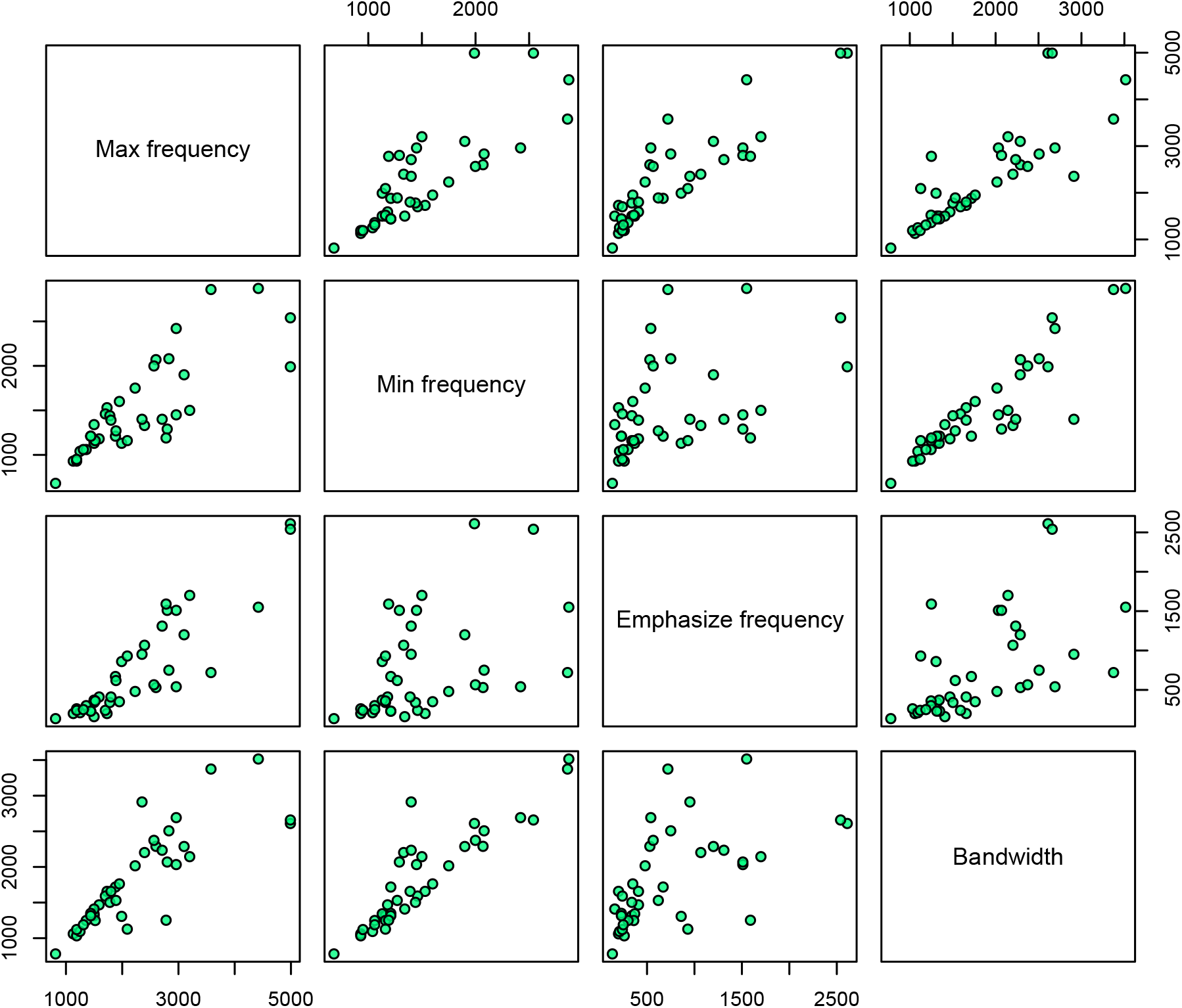
Scatterplots showing correlations among the four song variables.

